# Anoctamins mediate polymodal sensory perception and larval metamorphosis in a non-vertebrate chordate

**DOI:** 10.1101/2024.06.10.598279

**Authors:** Zonglai Liang, Jorgen Hoyer, Marios Chatzigeorgiou

## Abstract

An exceptional feature of most animals is their ability to perceive diverse sensory cues from the environment and integrate this information in their brain to yield ethologically relevant behavioral output. For the myriad of marine species, the ocean represents a complex sensory environment, which acts as a crucible of evolution for polymodal sensory perception. The cellular and molecular bases of polymodal sensory perception in a marine environment remain enigmatic.

Here we use Ca^2+^ imaging and quantitative behavioral analysis to show that in the tunicate *Ciona intestinalis* two members of the evolutionarily conserved Anoctamin family^1–4^ (Tmem16E/Ano5 and Tmem16F/Ano6), are required for sensing chemosensory and mechanosensory metamorphic cues. We find that they act by modulating neuronal excitability and Ca^2+^ response kinetics in the primary sensory neurons and axial columnar cells of the papillae, a widely conserved sensory-adhesive organ across ascidians^5–9^. Finally, we use electrophysiological recordings and a scramblase assay in tissue culture to demonstrate that Tmem16E/Ano5 acts as a channel, while Tmem16F/Ano6 is a bifunctional ion channel and phospholipid scramblase. Our results establish Ano5 and Ano6 as novel players in the zooplanktonic, pre-vertebrate molecular toolkit that controls polymodal sensory perception in aquatic environments.

## Results

The genome of tunicate Ciona intestinalis encodes four Tmem16/Ano proteins. We have recently demonstrated that Ciona Tmem16k/Ano10 functions in the developing notochord as an ion channel^10^. Our previous phylogentic analysis of the other three *Ciona* Anoctamins suggested that they belong to the Tmem16e/Ano5, Tmem16f/Ano6 and Tmem16g/Ano7 subfamilies^10^. Transcriptional fusion analysis showed that both Tmem16e/Ano5 and Tmem16f/Ano6 (here after referred to as Ano5 and Ano6 respectively) are expressed primarily in the central and peripheral nervous system of the larvae (Figure 1A, 1B; Figure S1A-D). Ano5 and Ano6 were expressed in the primary sensory neurons (PSNs) and axial columnar cells (ACCs) of the sensory adhesive structures called papillae which mediate larval settlement and the onset of metamorphosis in ascidians^5,9,11–15^ (Figure 1A, 1B; Figure S1A-G). To determine the subcellular localization of Ano5 and Ano6 generated translational fusions (Figure 1C). We found that Ano5 localized primarily to the ER but not the cilia and digitiform projections of the PSNs and ACCs respectively. In contrast, Ano6 localized to the plasma membrane of the cell bodies and was enriched in the cilia and digitiform projections of the PSNs and ACCs.

**Figure 1.**
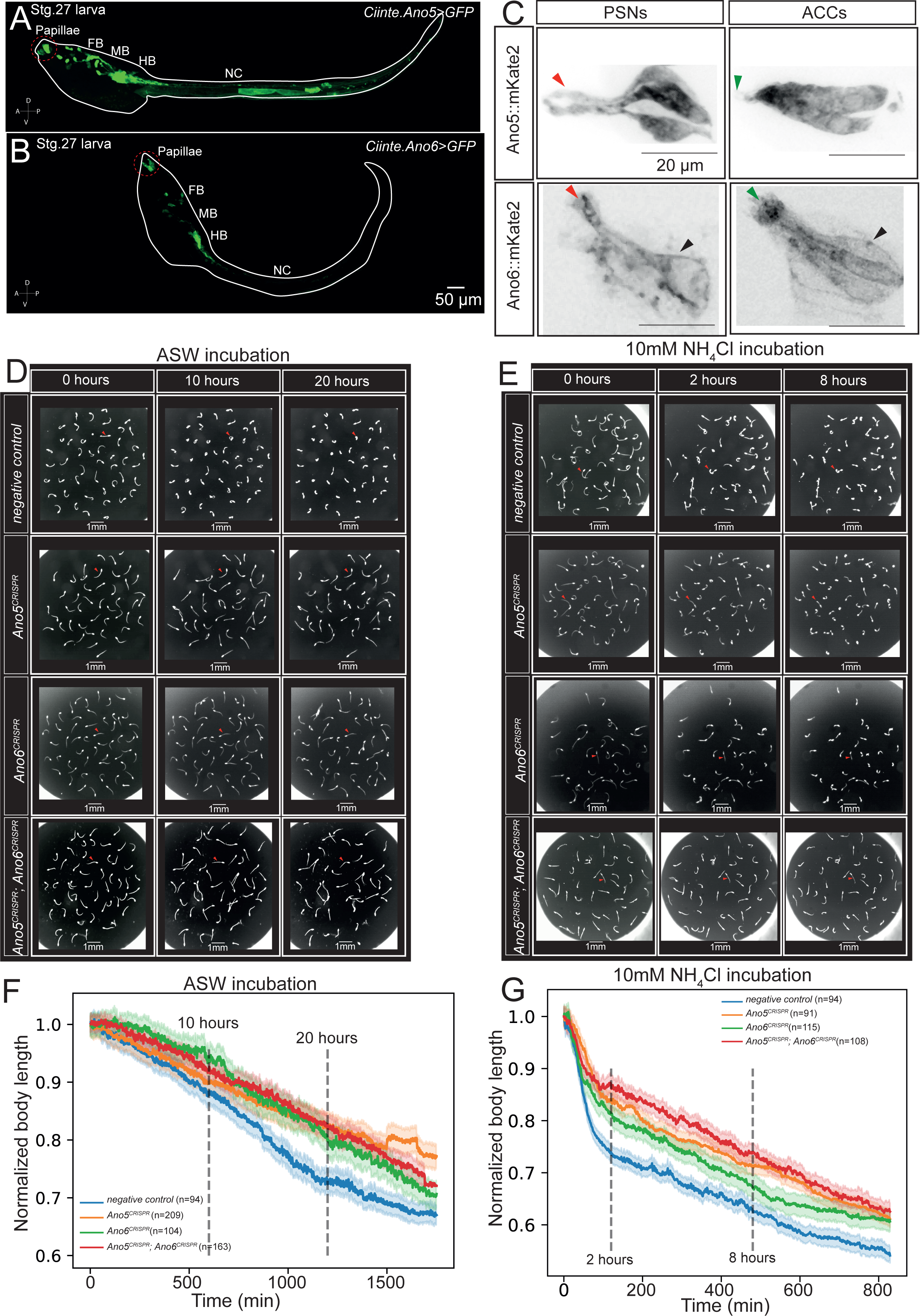
Ano5 and Ano6 localize to the sensory endings and the ER of the papillae and regulate the rates of larval metamorphosis. (A-B) Example maximal projections of confocal stacks showing the expression of transgenic larvae expressing (A) *Ciinte.Ano5>GFP* and (B) *Ciinte.Ano6>GFP*. Scale bar is 50μm. Abbreviations: FB= Forebrain, MB=Midbrain, HB=Hindbrain, NC=Nerve cord. Red dashed circles highlight the papillae. (C) Confocal micrographs of translational fusions of Ano5::mKate2 and Ano6::mKate2 drive by either the *Ciinte.pc2* promoter which drives expression in the PSNs or the *Ciinte.βγ-crystallin* promoter which drives expression in the ACCs. Red and green arrowheads point to the ciliary sensory ends of PSNs and the sensory protrusions of the ACCs respectively. (D) Example video frames from negative control, Ano5^CRISPR^, Ano6^CRISPR^ and Ano5^CRISPR^; Ano6^CRISPR^ larvae at three different time points: 0 hours, 10 hours, and 20 hours in the presence of ASW. (E) Example video frames for the same genotypes at three different time points: 0 hours, 2 hours, and 8 hours in the presence of 10mM NH_4_Cl which is a metamorphosis promoting cue. Red arrowheads point to example larvae in the process of tail regression. (F, G) Normalized body length curves for negative control, Ano5^CRISPR^, Ano6^CRISPR^ and Ano5^CRISPR^; Ano6^CRISPR^ larvae in the presence of (F) ASW or (G) 10mM NH_4_Cl. Solid lines: mean; shaded areas: standard error of the mean (SEM). Number of animals used are indicated in the brackets. For further quantification of normalized body length at different time points please see Figure S2A-D and Data S1B-S1E.

**Figure S1.**
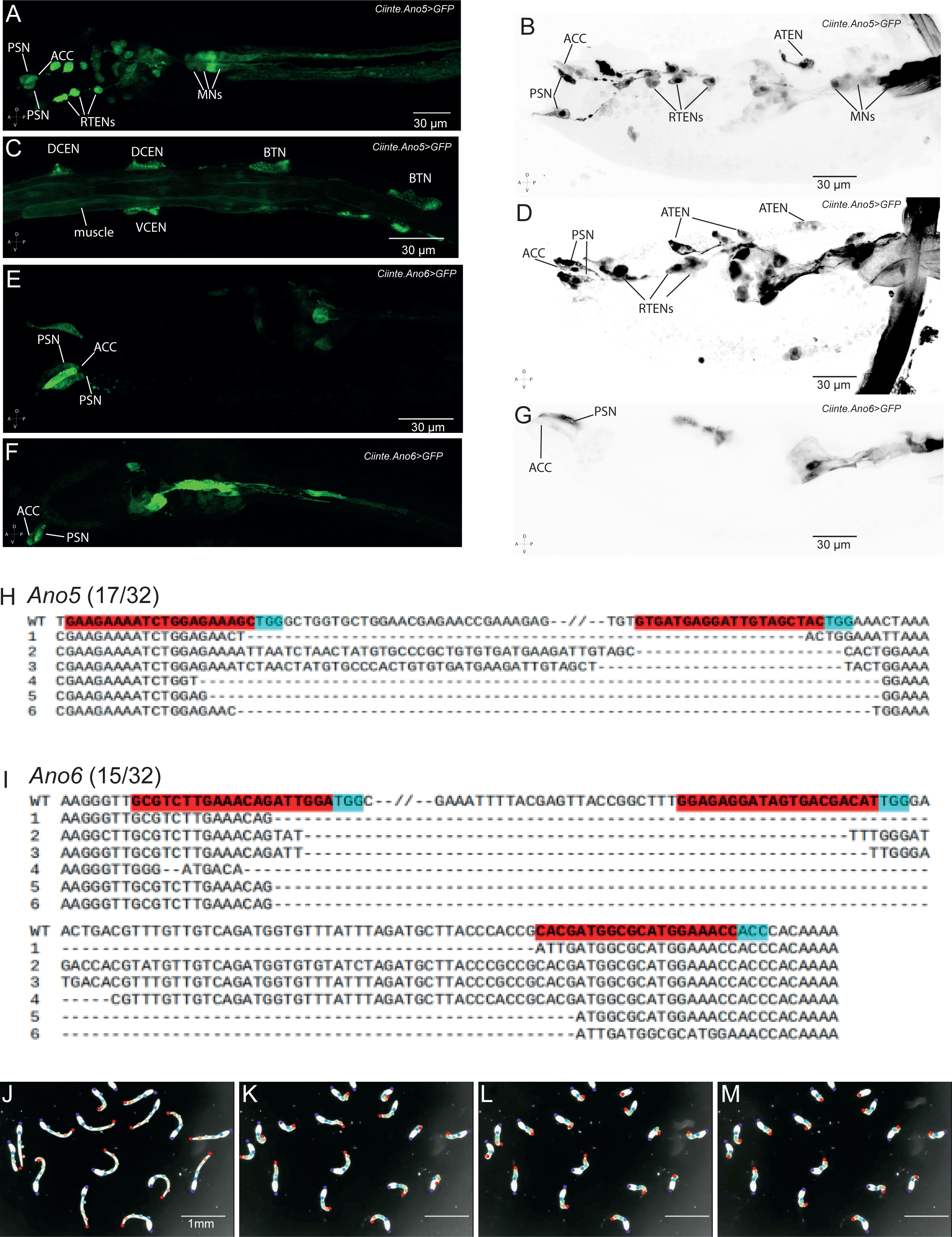
Examples of reporter expression, genome editing results and markerless pose estimation for Ano5 and Ano6. Related to Figure 1. (A-D) Example maximal projections of confocal stacks showing the expression of transgenic larvae expressing *Ciinte.Ano5>GFP* and (E-G) Example maximal projections of confocal stacks showing the expression of transgenic larvae expressing *Ciinte.Ano6>GFP*. Scale bars are 30μm. Abbreviations: primary sensory neurons (PSNs), axial columnar cells (ACCs), rostral trunk epidermal neurons (RTENs), Motorneurons (MNs), apical trunk epidermal neurons called (ATENs), dorsal caudal epidermal neuron (DCENs), ventral caudal epidermal neuron (VCENs), bipolar tail neuron (BTNs). Anterior to the left. (H-I) Sequencing detection of the deletions induced by CRISPR/Cas9 of the (H) *Ano5* and (I) *Ano6* loci. The number of clones with deletions are indicated (17/32 and 15/32 for *Ano5* and *Ano6* respectively). gRNA target sequence is highlighted in red, while the PAM sequence is highlighted in cyan. area, cyan-color labels the PAM sequence. (J-M) Example video frames from a metamorphosis video where the DLC points are shown on each animal. Scale bar is 1mm

### Ano5 and Ano6 are required for metamorphosis induced by mechanical and chemical cues

The papillae mediate ascidian larval settlement and metamorphosis in response to a diversity of mechanical and chemical cues^5^. During this process the larvae will use their papillae to identify a suitable substrate to which they will adhere by means of secretion of adhesives by the papillar collocyte cells. This process is followed by the gradual regression of the tail into the larval trunk^11^. We wondered whether Ano5 and Ano6 are required for metamorphosis. To address this question, we leveraged CRISPR/Cas9 technology to generate *Ano5^CRISPR^*and *Ano6^CRISPR^* single and double mutants, which we assayed in a larval metamorphosis assay combining long-term, continuous animal tracking^16^ with markerless pose estimation^17^ (Figure 1D-G; Figure S1J-M). Initially we compared the ability of the larvae to undergo metamorphosis in artificial sea water, where the only stimulus was the presence of a mechanical cue coming from the stiff plastic dish tracking arena (Figure 1D). Negative control larvae progressively regressed their tail over a period of 29 hours, as the normalized body length curve demonstrates (Figure 1F). We compared the normalized body length of larvae between negative control, *Ano5^CRISPR^* and *Ano6^CRISPR^*single and double mutants at two different time points (10, 20 hours). At both the 1^st^ and the 2^nd^ time point all three mutant genotypes showed a significant difference compared to negative controls (Figure 1F; Figure S2A, S2B; Data S1B, S1C).

We repeated the experiment, this time presenting the larvae with 10mM NH_4_Cl, a chemical stimulus that is known to promote metamorphosis^5^ (Figure 1E, 1G). The combined presence of a mechanical stimulus together with 10mM NH_4_Cl accelerated (>2x faster compared to ASW alone) metamorphosis rates confirming previous observations^5^ (Figure 1E, 1G). We compared the normalized body length of the larvae 2 hours and 8 hours after the onset of the assay across all the genotypes (Figure 1E, 1G; Figure S2C, S2D; Data S1D, S1E). All mutant genotypes were significantly slower to metamorphose compared to negative controls after 2 hours of incubation with 10mM NH_4_Cl. However, at the 8 hour time-point, *Ano6^CRISPR^* animals caught-up with negative controls in contrast to *Ano5^CRISPR^*and *Ano5^CRISPR^; Ano6^CRISPR^* double mutants (Figure 1E, 1G; Figure S2C, S2D; Data S1D, S1E). Taken together our experiments show that Ano5 and Ano6 modulate the rate of metamorphosis in the presence of mechanical and chemical cues.

### Ano5 and Ano6 control the Ca^2+^ response frequency and kinetics of the PSNs and ACCs to mechanical and chemical stimuli

Recently we demonstrated that the PSNs and ACCs are polymodal cells capable of eliciting Ca^2+^ responses to diverse mechanical and chemical stimuli that promote or impede settlement and metamorphosis^5^. Since Ano5 and Ano6 are expressed in the PSNs and ACCs we wondered whether *Ano5^CRISPR^*and *Ano6^CRISPR^* mutants affected the frequency of Ca^2+^ responses and the features of the Ca^2+^ transients in these cell types in response to mechanical and chemical stimuli (Figure 2A-L; Figure S2E-X).

To examine the function of *Ano5^CRISPR^* and *Ano6^CRISPR^* single or double mutants in the PSNs we used the *Ciinte.pc2* promoter to drive the expression of the genetically encoded calcium indicator (GECI) GCaMP6s. Using a piezo driven glass probe we were able to deliver mechanical poke stimuli to the papillae, in response to which we obtained Ca^2+^ transients from the PSNs of negative control and mutant larvae (Figure 2A, 2C, 2D; Figure S2E-S2I; Data S1F-S1J). The frequency of Ca^2+^ responses in *Ano6^CRISPR^*mutants was reduced relative to negative controls. Both Ano5 and Ano6 contribute to the Ca^2+^ transient characteristics. In particular *Ano5^CRISPR^*mutants were characterized by a significantly higher peak amplitude and area compared to negative controls, while Ca^2+^ transients in *Ano6^CRISPR^* animals exhibited significantly longer peak rise times (Figure 2A, 2C; Figure S2E-S2I; Data S1F-S1J). *Ano5^CRISPR^*; *Ano6^CRISPR^* double mutants showed Ca^2+^ responses to poking characterized by longer peak rise period compared to negative controls (Figure 2A, 2C; Figure S2E-S2I; Data S1F-S1J). *Ano5^CRISPR^*; *Ano6^CRISPR^* responses were significanlty different from *Ano5^CRISPR^*single mutants but indistinguishable from *Ano6^CRISPR^* mutants (Figure 2A, 2C; Figure S2E-S2I; Data S1F-S1J).

**Figure 2.**
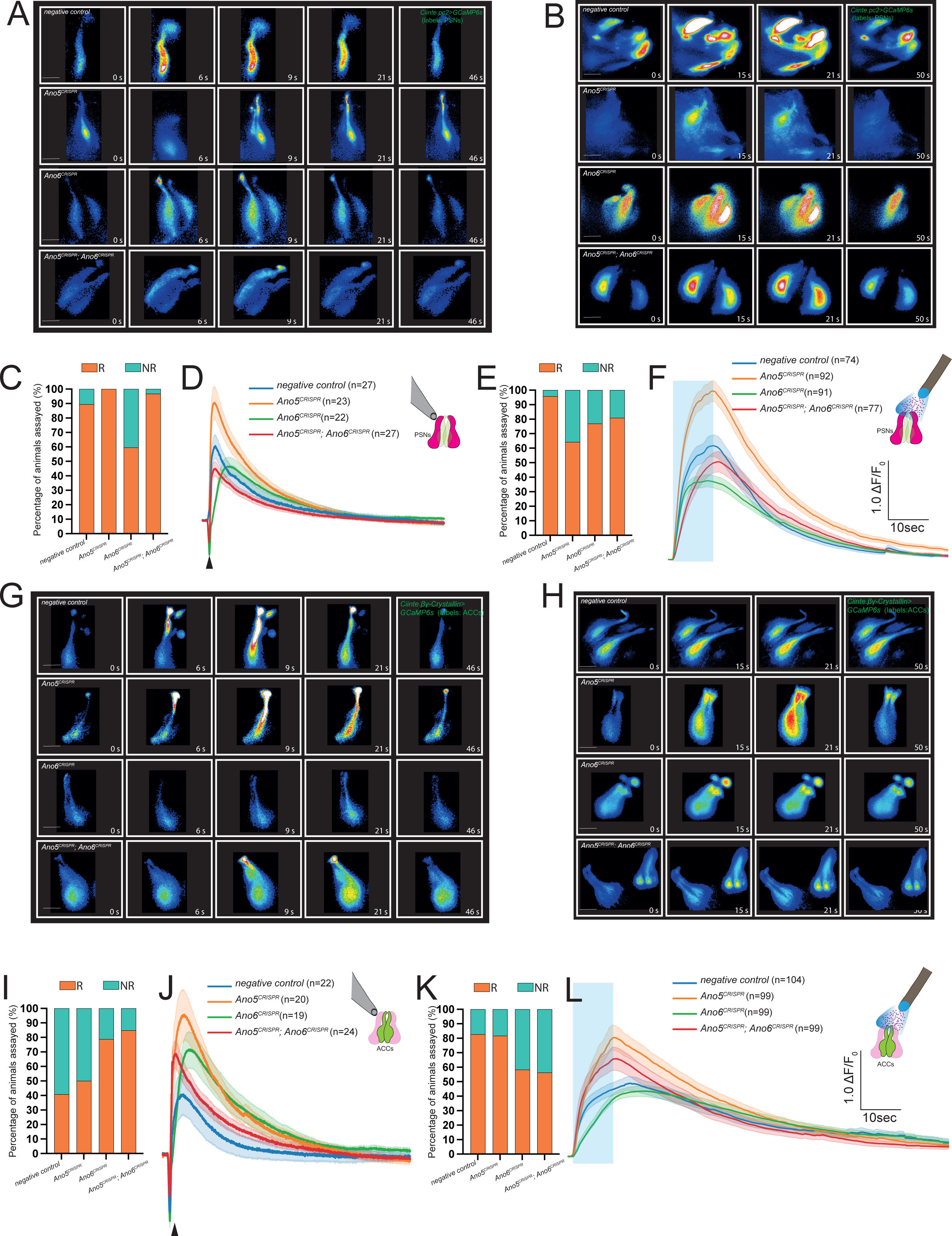
Ano5 and Ano6 modulate calcium transients in PSNs and ACCs in response to mechanical and chemical stimuli. (A, B) Montage of PSN Ca^2+^ responses to (A) mechanical poke and (B) 10mM NH_4_Cl from negative control, Ano5^CRISPR^, Ano6^CRISPR^ and Ano5^CRISPR^; Ano6^CRISPR^ larvae expressing the GECI GCaMP6s under the *Ciinte.pc2* promoter. We presented the mechanical stimulus on the 5^th^ second of the recording and the chemical stimulus between the 10^th^ and 20^th^ seconds. Scale bar= 20μm (C) Quantification of the fraction of animals with PSNs which respond to mechanical poke (R=Responders; NR=Non-Responders). (D) Average traces of Ca^2+^ responses from animals responding to mechanical poke across all four genotypes assayed. Solid lines indicate the mean trace and shaded areas correspond to SEM. Number of traces contributing to the average traces are shown in parentheses. (E) Quantification of the fraction of animals with responsive PSNs to 10mM NH_4_Cl stimulation across the four genotypes. (F) Average traces of Ca^2+^ responses from animals responding to 10mM NH_4_Cl stimulation across all four genotypes assayed. Blue bar indicates the period of stimulation. (G-H) Montage of ACC Ca^2+^ responses to (G) mechanical poke and (H) 10mM NH_4_Cl from negative control, Ano5^CRISPR^, Ano6^CRISPR^ and Ano5^CRISPR^; Ano6^CRISPR^ larvae expressing the GECI GCaMP6s under the *Ciinte.βγ-crystallin* promoter. Scale bar= 20μm. (I) Quantification of the fraction of animals with ACCs which respond to mechanical poke (R=Responders; NR=Non-Responders). (K) Quantification of the fraction of animals with responsive ACCs to 10mM NH_4_Cl stimulation across the four genotypes. (L) Average traces of Ca^2+^ responses in response to 10mM NH_4_Cl across all four genotypes. Blue bar indicates the 10 second period of stimulation with 10mM NH_4_Cl. Further quantification of peak features can be found in Figure S2E-X and Data S1F-S1Y

**Figure S2.**
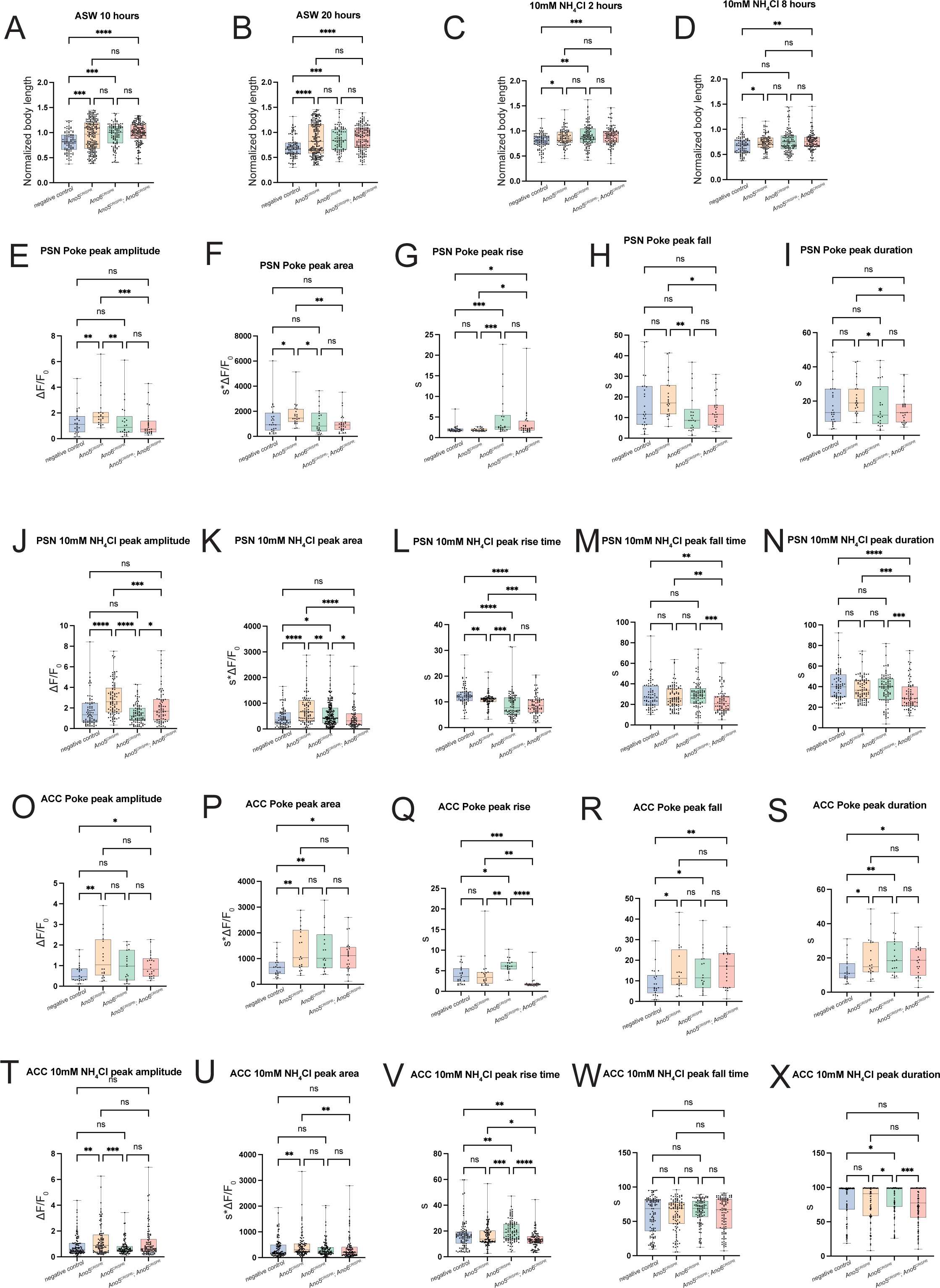
Quantification of metamorphosis behavioral parameters and calcium peak features across all for genotypes. Related to Figure 2. (A, B) Quantification of normalized body length across four genotypes (negative control, Ano5^CRISPR^, Ano6^CRISPR^ and Ano5^CRISPR^; Ano6^CRISPR^) in ASW at 10 hours (panel A) and 20 hours (panel B) after the start of the assay. (C, D) Box plots of normalized body length across four genotypes in the presence of 10mM NH_4_Cl at 2 hours (panel C) and 8 hours (panel B) after the start of the assay. For statistical analysis see Data S1B-S1E. (E-X) Box plots quantifying PSN and ACC Ca^2+^ transient peak amplitude, peak area, peak rise time, peak fall time and peak duration features across all four genotypes in (E-I) PSNs stimulated with a mechanical poke, (J-N) PSNs stimulated with 10mM NH_4_Cl, (O-S) ACCs stimulated with a mechanical poke and (T-X) ACCs stimulated with 10mM NH_4_Cl. For statistical analysis please see Data S1F-S1Y.

Subsequently, we tested the ability of *Ano5^CRISPR^* and *Ano6^CRISPR^* single and double mutants to affect the Ca^2+^ responses of the PSNs to a chemical stimulus. Negative controls responded robustly to 10mM NH_4_Cl as previously reported^5^ (Figure 2B, 2D, Figure S2J-S2N; Data S1K-S1O). *Ano5^CRISPR^* Ca^2+^ responses 10mM NH_4_Cl were characterized from a significantly faster rising time, higher peak amplitude and larger area under the peak compared to negative controls (Figure 2B, 2D, Figure S2J-S2N; Data S1K-S1O). The Ca^2+^ transients in of *Ano6^CRISPR^* animals exhibited significantly smaller peak area and very short peak rise time compared to negative controls controls (Figure 2B, 2D, Figure S2J-S2N; Data S1K-S1O). Most of the double mutant Ca^2+^ peak features were significantly different relative to *Ano5^CRISPR^*and *Ano6^CRISPR^* single mutants, with values closer to those of negative control Ca^2+^ peak features suggesting that Ano5 and Ano6 have a supresive function in the context of PSN chemosensation (Figure 2B, 2D, Figure S2J-S2N; Data S1K-S1O).

To examine the function of *Ano5^CRISPR^* and *Ano6^CRISPR^* single or double mutants in the ACCs we used the *Ciinte.βγ-crystallin* promoter to drive the expression of the genetically encoded calcium indicator (GECI) GCaMP6s (Figure 2G, 2H). Negative controls elicited elicited Ca^2+^ transients in response to mechanical stimulation in ∼40% of the stimulation trials (Figure 2I). The ACCs of *Ano5^CRISPR^* and *Ano6^CRISPR^* single and double mutant larvae showed increased response frequency when stimulated with a mechanical stimulus compared to negative control. This suggests that Ano5 and Ano6 attenuate the sensitivity of the ACCs to mechanical cues. Loss of Ano5 elicited Ca^2+^ transients with signficantly higher peak amplitude, area and fall time relative to negative controls (Figure 2J; Figure S2O-S2S; Data S1P-T). Loss of Ano6 led to an increase in magntitude of all Ca^2+^ peak features other than peak amplitude (Figure 2J; Figure S2O-S2S; Data S1P-T). Finally, *Ano5^CRISPR^*; *Ano6^CRISPR^*double mutants all peak features that we quantified were significantly different to negative controls. In contrast only the peak rise time was signficantly different between either of the single mutants and the double mutant (Figure 2J; Figure S2O-S2S; Data S1P-T).

Finally, we tested the ability of the ACCs to respond to 10mM NH_4_Cl in negative control and mutant backgrounds. Consistent with previous observations the ACCs of negative control larvae responded to ∼82% of the trials with 10mM NH_4_Cl (Figure 2H, 2K, 2L). The ACCs of *Ano5^CRISPR^* mutants responded with a similar frequency to controls (∼81.7%) in contrast to *Ano6^CRISPR^* and double mutants which responded less frequently to mechanical stimulation trials (∼58% and ∼56% respectively) (Figure 2H, 2K, 2L). Ammonia evoked Ca^2+^ transients in the ACCs of *Ano5^CRISPR^*larvae showed significantly higher peak amplitude and area compared to negative controls (Figure 2L; Figure S2T-X; Data S1U-S1Y). In contrast, Ca^2+^ transients from *Ano6^CRISPR^* larvae were almost indistinguishable from negative controls aside from a significantly slower peak rise time (Figure 2L; Figure S2T-X; Data S1U-S1Y). Ca^2+^ responses of *Ano5^CRISPR^*; *Ano6^CRISPR^* double mutant ACCs were characterized by faster peak rise times. showed Ca^2+^ responses that were larger than those of negative controls but this phenotype was not statistically significant with the exception of peak rise time (Figure 2L; Figure S2T-X; Data S1U-S1Y).

### Ano5 and Ano6 exhibit channel properties but Ano6 acts also as a phospholipid scramblase

Previous studies have classified TMEM16/ANO family members as CaCCs and Ca^2+^-dependent phospholipid scramblases, while some TMEM16/ANO phosopholipid scramblases have also been shown to conduct ions (reviewed in^3,4,18^). To investigate whether *Ciona* Ano5 and Ano6 act as channels or phospholipid scramblases, we performed heterologous expression experiments. First we tested whether Ano5 and Ano6 act as phospholipid scramblases. We found that Ano5 transfected cells showed a limited ability (4.24% +ve cells) to cause Phosphatidyl-Serine exposure upon stimulation with the Ca^2+^ ionophore ionomycin (Figure 3A; Data S1Z). In contrast, Ano6

**Figure 3.**
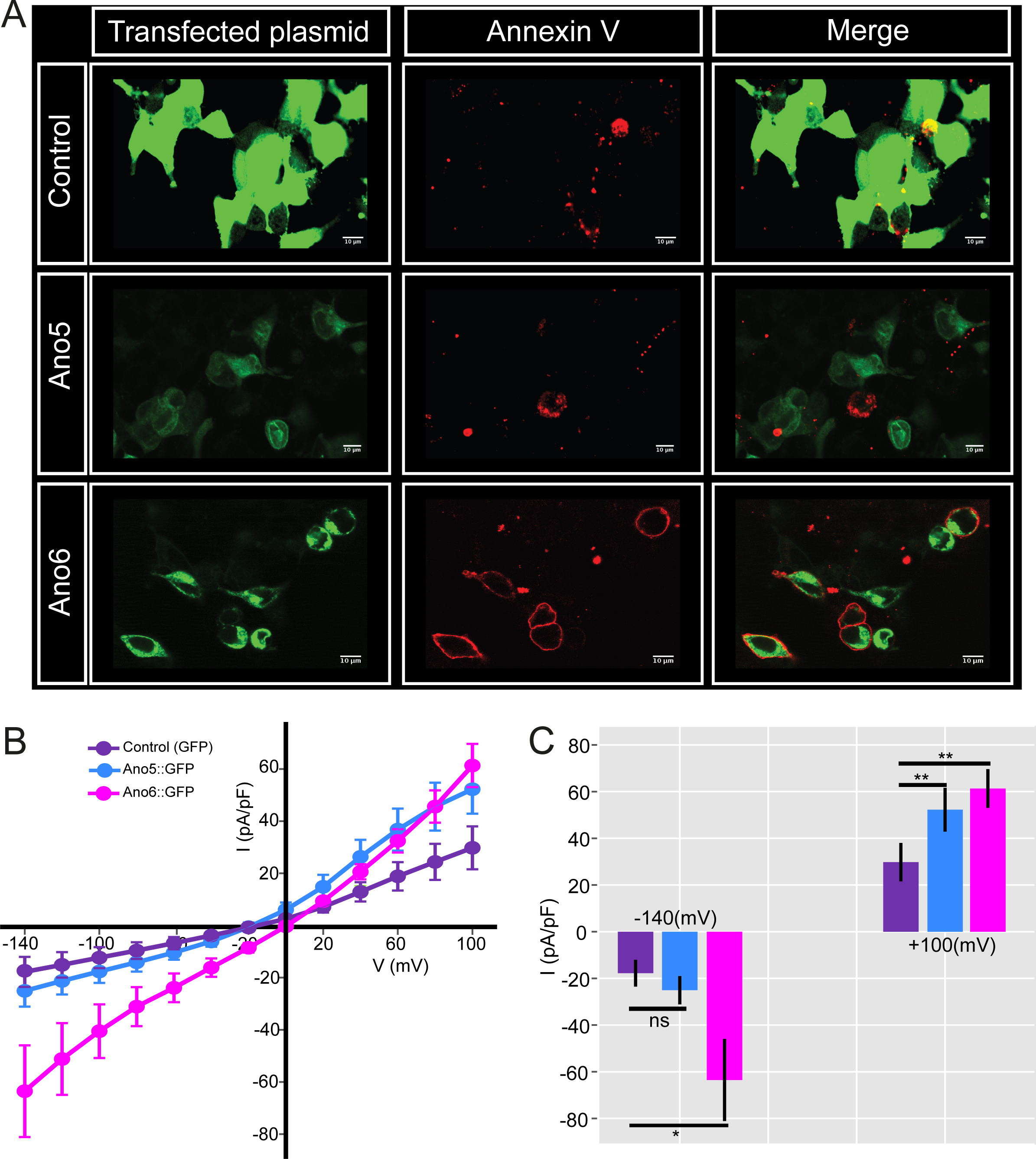
Heterologous expression of Ano5 and Ano6 reveals that both exhibit channel properties but Ano6 acts also as a phospholipid scramblase. (A) Scramblase assay in HEK293T cells overexpressing control (soluble GFP), Ano5::GFP and Ano6::GFP stimulated with ionomycin. GFP signal (green), Alexa Fluor™ 568 conjugated Annexin V signal (red). Scale bars= 10μm. See Data S1Z for quantification. (B) Stimulation with 100μM ATP activates currents in patch-clamp experiments. HEK293T cells were held at 0 mV and pulsed to voltages from −140mV to +100mV (in steps of 20mV). Corresponding current/voltage relationships for control cytoplasmic GFP (n=13), Ano5::GFP (n=20) and Ano6::GFP (n=17). (C) Quantification of the currents from B at −140mV and +100 mV. transfected cells showed significant scramblase activity upon treatment with ionomycin (13.02% +ve cells) when compared to mock transfected cells stimulated with ionomycin (2.25% +ve cells) (Figure 3A, Data S1Z).

Finally, we employed electrophysiology to determine whether Ano5 and Ano6 possess an ion channel functionality. We co-expressed Ano5 or Ano6 and the purinergic receptor P2Y2R (which is coupled to Gq signaling and leads to an increase in intracellular Ca^2+^ upon stimulation by ATP) in HEK293T cells and tested for current activity in response to 10μM ATP stimulation in whole-cell configuration. Both Ano5 and Ano6 transfected cells exhibited larger currents compared to mock transfected cells (Figure 3B, 3C). Ano5 exhibited outwardly rectifying currents, while Ano6 showed a rather linear current/voltage relationship. Our experimental evidence argues in favor of Ano5 working as a channel and Ano6 working as both a channel and phospholipid scramblase.

## Discussion

This study establishes *Ciona* Ano5 and Ano6 as two novel members of the molecular toolkit underlying polymodal sensory transduction in marine planktonic larvae.

We demonstrate that Ano5 and Ano6 are expressed in the nervous system of *Ciona* larvae including the PSNs and ACCs of the papillar organs. These are polymodal cells capable of sensing mechanical as well as gustatory and olfactory chemical cues^5^. The papillae develop from a region anterior to the neural plate, which has been postulated to be an olfactory/adenohypophyseal placode homologue, expressing several regulatory genes that are required for olfactory placode development in vertebrates^19–26^. Interestingly, the mammalian TMEM16B/ANO2, TMEM16J/ANO9 proteins are expressed in olfactory sensory neurons (OSNs) and contribute to signaling in the olfactory epithelium^27–31^. This suggests that the role of TMEM/ANO proteins in olfaction emerged in early chordates, and it has been conserved in vertebrates.

To date our understanding of the sensory transduction machinery underlying larval metamorphosis is limited. A previous study had reported that the TRP channel PKD2 is required for the onset of metamorphosis in response to mechanical input^32^. Here we we have shown that both Ano5 and Ano6 are required to achieve robust tail regression in the presence of mechanical and chemical metamorphic cues. However, given that loss of Ano5 and Ano6 does not lead to complete inhibition of metamorphosis it is reasonable to hypothesize that additional unknown sensory transduction molecules modulate larval metamorphosis.

At the cellular level we found that Ano5 and Ano6 control the frequency with which the PSNs and ACCs respond to mechanical and chemical stimuli. At the same time, they shape the Ca^2+^ response kinetics. Specifically, Ano5 ‘s primary function is to dampen Ca^2+^ responses elicited by the PSNs and the ACCs in response to both mechanical and chemical stimulation. We also found that it is selectively involved in reducing the frequency with which the PSNs respond to 10mM NH4Cl. Recent studies highlight the role of mammalian TMEM16/ANO proteins in the modulation of neuronal excitability as they can either increase or decrease excitability^33–38^. Several studies have reported that loss of TMEM16B/ANO2 can increase spiking activity in OSNs, while leading to olfactory behavioral deficits ^33,38^. Similarly to mammalian TMEM16B/ANO2, *Ciona* Ano5 may participate in a sensory transduction cascade where it serves as part of a feedback mechanism to ‘clamp’ PSN and ACC output to the downstream circuit.

On the other hand, we determined that Ano6 plays an important role during the onset phase kinetics of the Ca^2+^ response to both mechanical and chemical stimuli. Loss of Ano6 slows down the rising phase of the Ca^2+^ transients in both the PSNs and ACCs during mechanical and chemical stimulation. The impact of the amplitude and kinetics of sensory transduction to multimodal-driven behaviors is not very well understood, partly due to the lack of experimental paradigms where this phenomenon can be explored.

The genetic interactions we explored via the *Ano5^CRISPR^*; *Ano6^CRISPR^* double mutant suggest that the frequency of the responses and the response kinetics of the double mutants look more like those of *Ano6^CRISPR^* animals. This suggests that Ano6 exerts an epistatic effect on Ano5 activity. Interestingly, TMEM16B/ANO2 and TMEM16F/ANO6 have been shown to be co-expressed and even form heteromeric interactions in olfactory sensory neurons of mice^39^. Future studies will shed light on whether Ano5 and Ano6 establish heteromeric interactions in the PSN and ACC cells.

Here, we have shown that Ano5 functions as a channel, while Ano6 is able to act as a phospholipid scramblase and as a channel. Previously examples of such bifunctional TMEM16/ANO proteins include mammalian TMEM16F/ANO6^40–43^, TMEM16K/ANO10^44,45^ and the fungal homologs afTMEM16^46^ and nhTMEM16^47,48^. Our data suggest that the presence of bifunctional TMEM16F/ANO6s is conserved in the chordate lineage and that this bifunctionality of TMEM16F/ANO6 has been preserved following the expansion of the TMEM16/ANO family in vertebrates.

Taken together our work raises the possibility that the adoption of the TMEM16/Ano family to the molecular toolkit mediating olfaction arose at the base of chordates.

## KEY RESOURCES TABLE

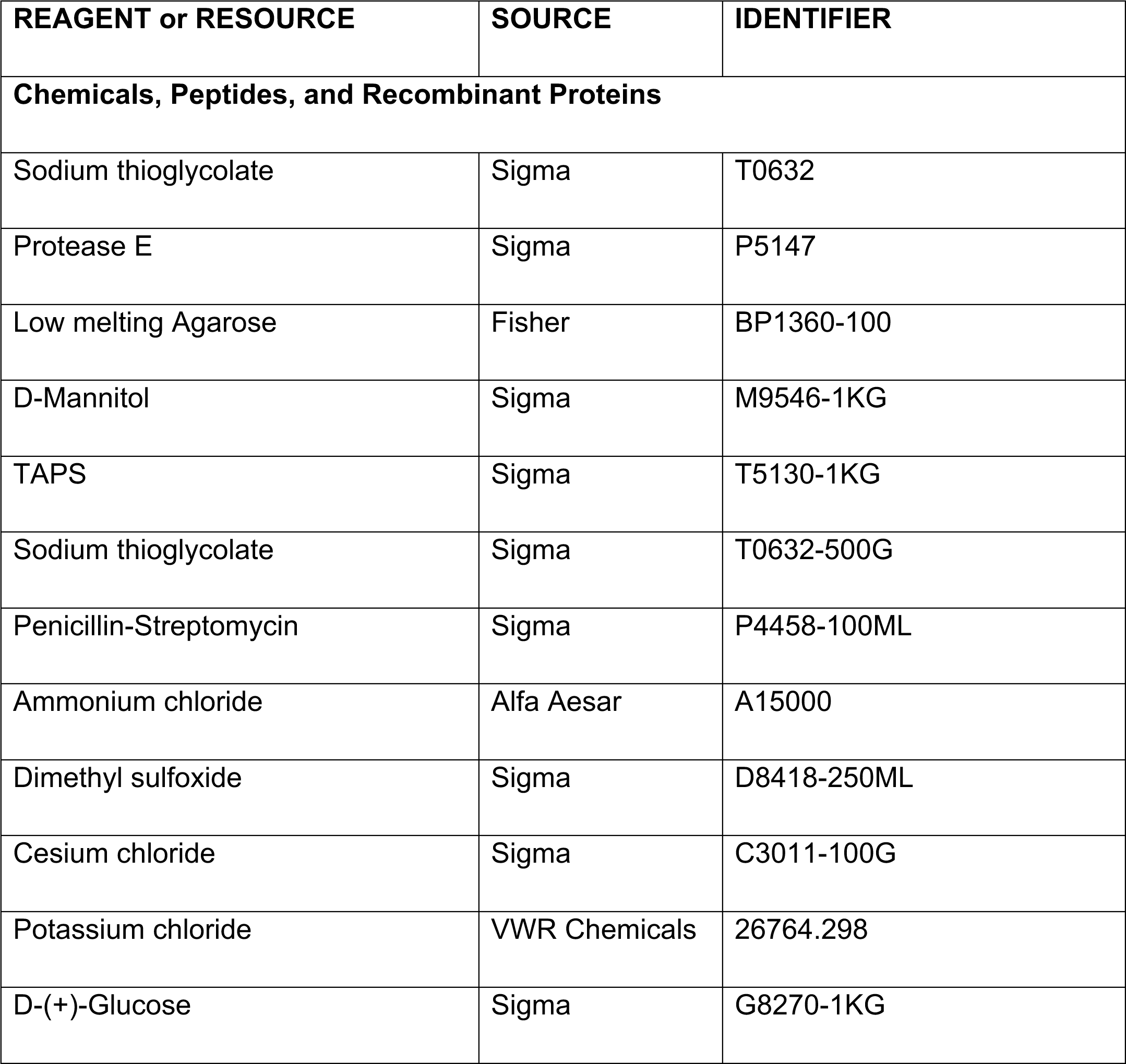
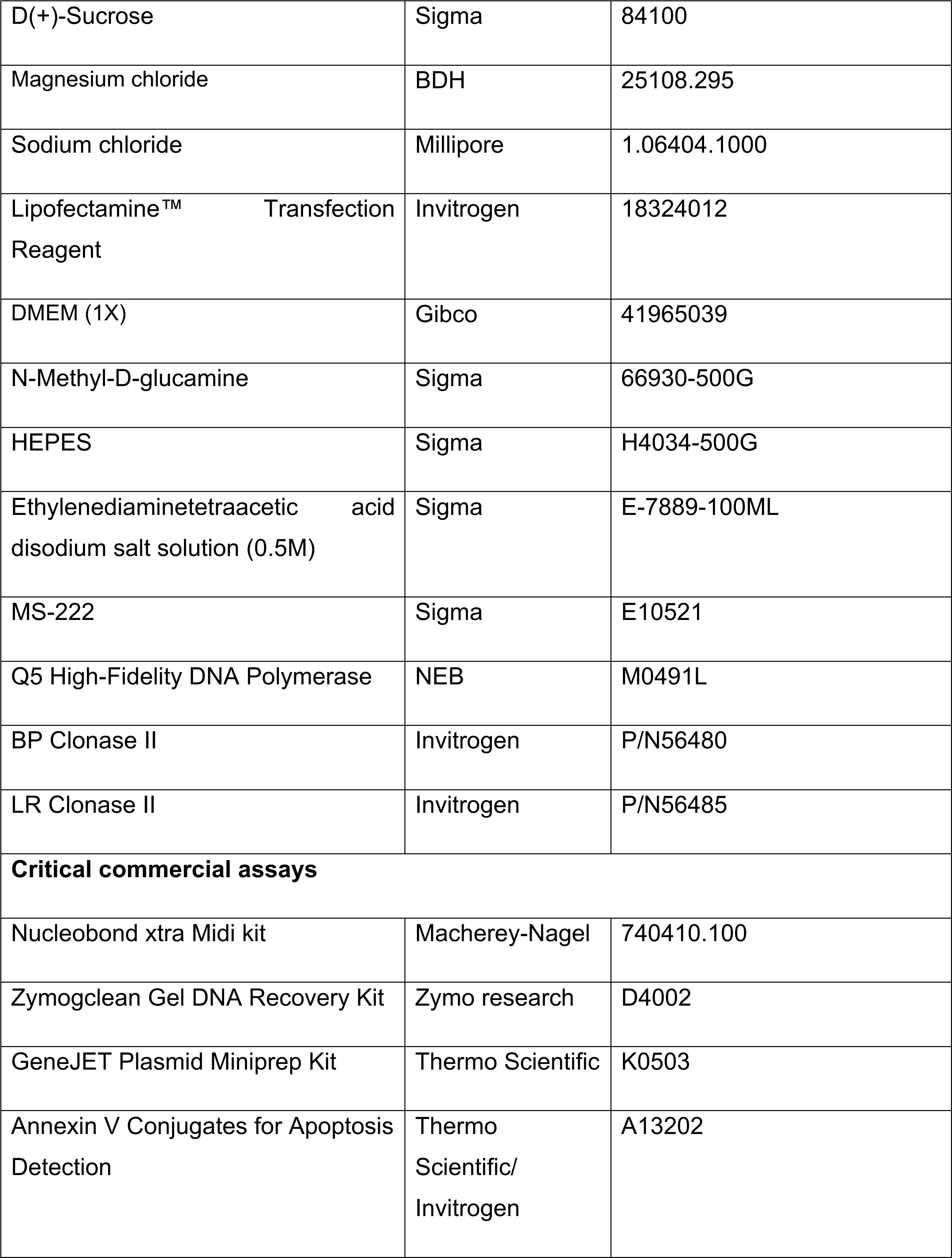
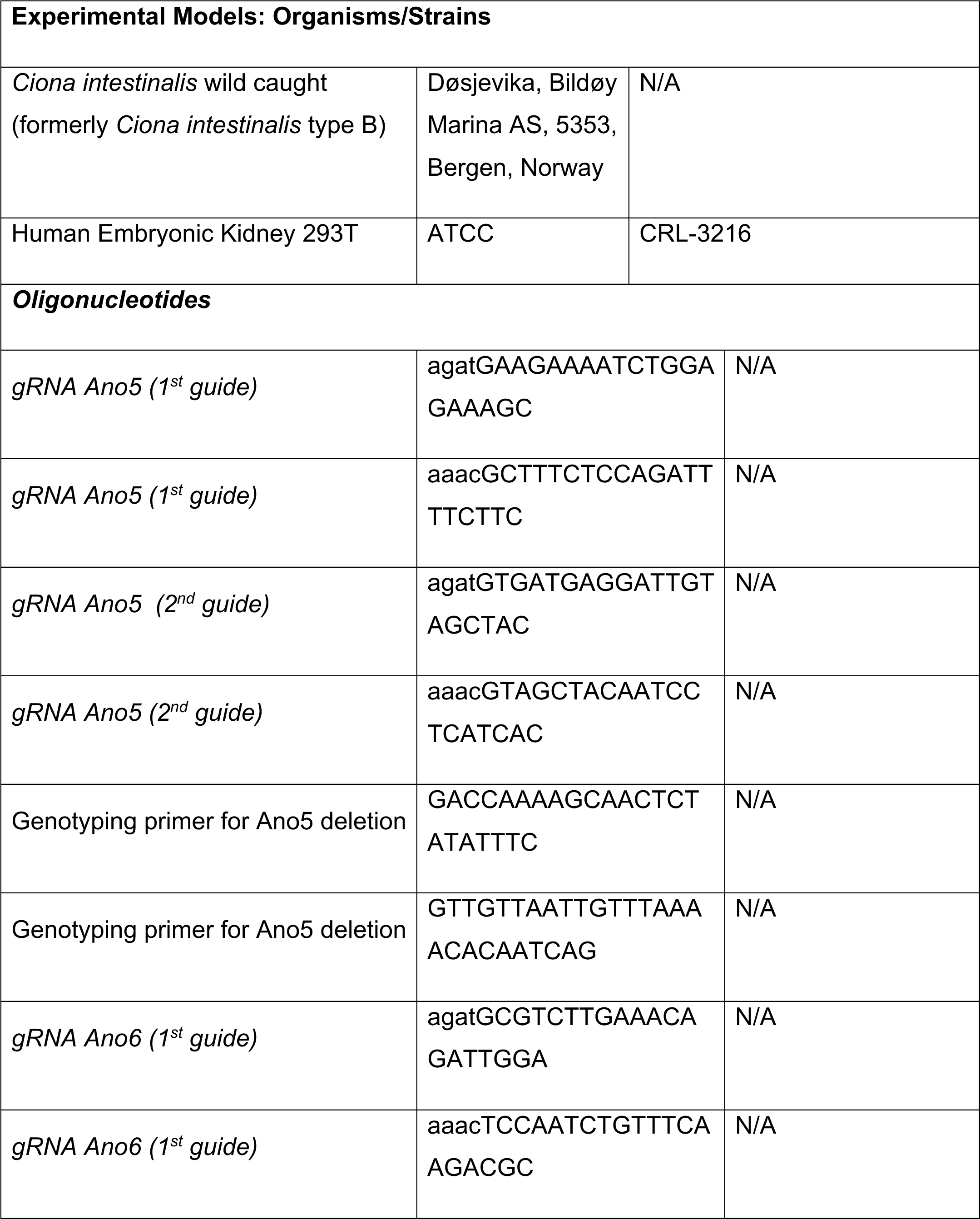
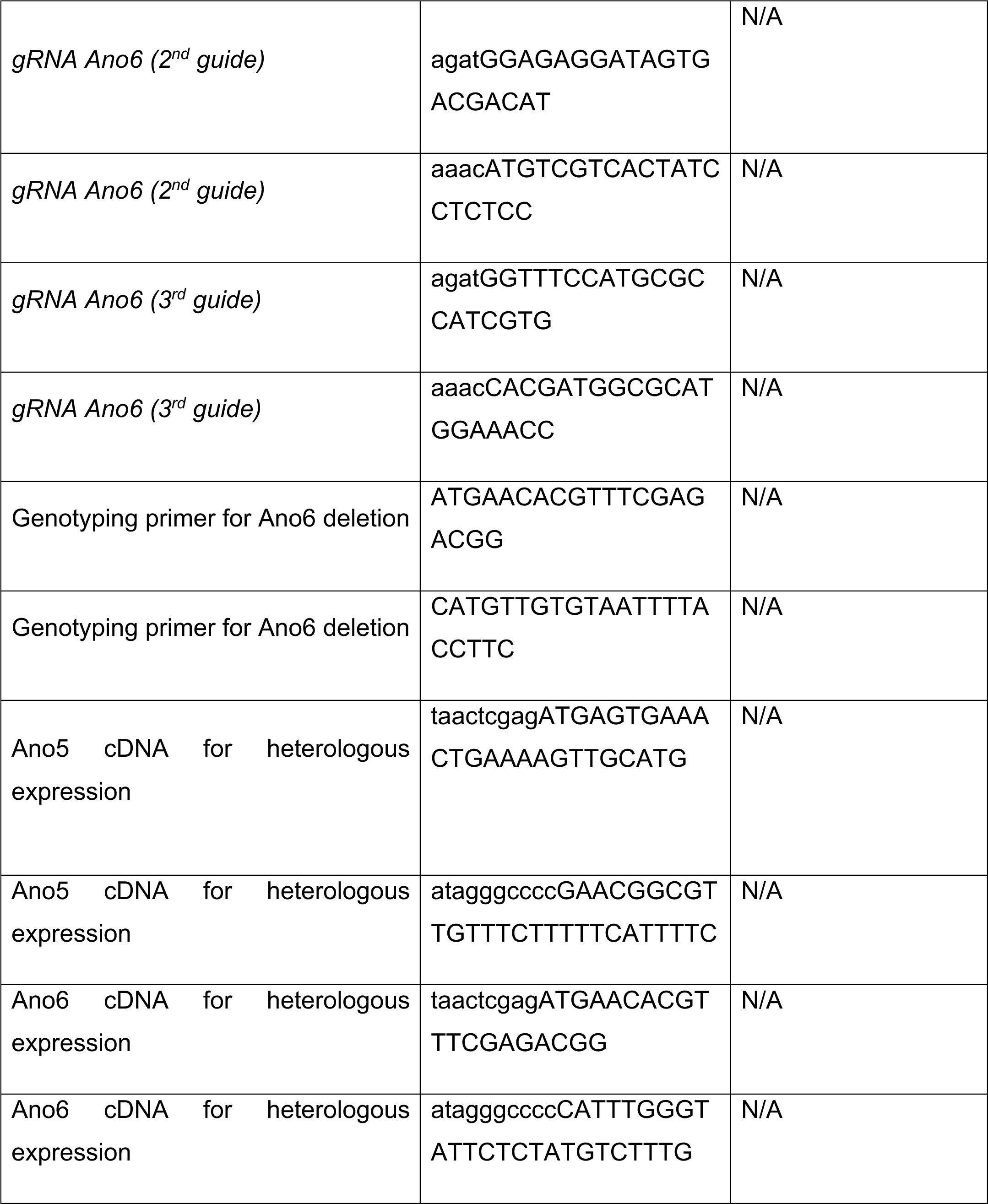
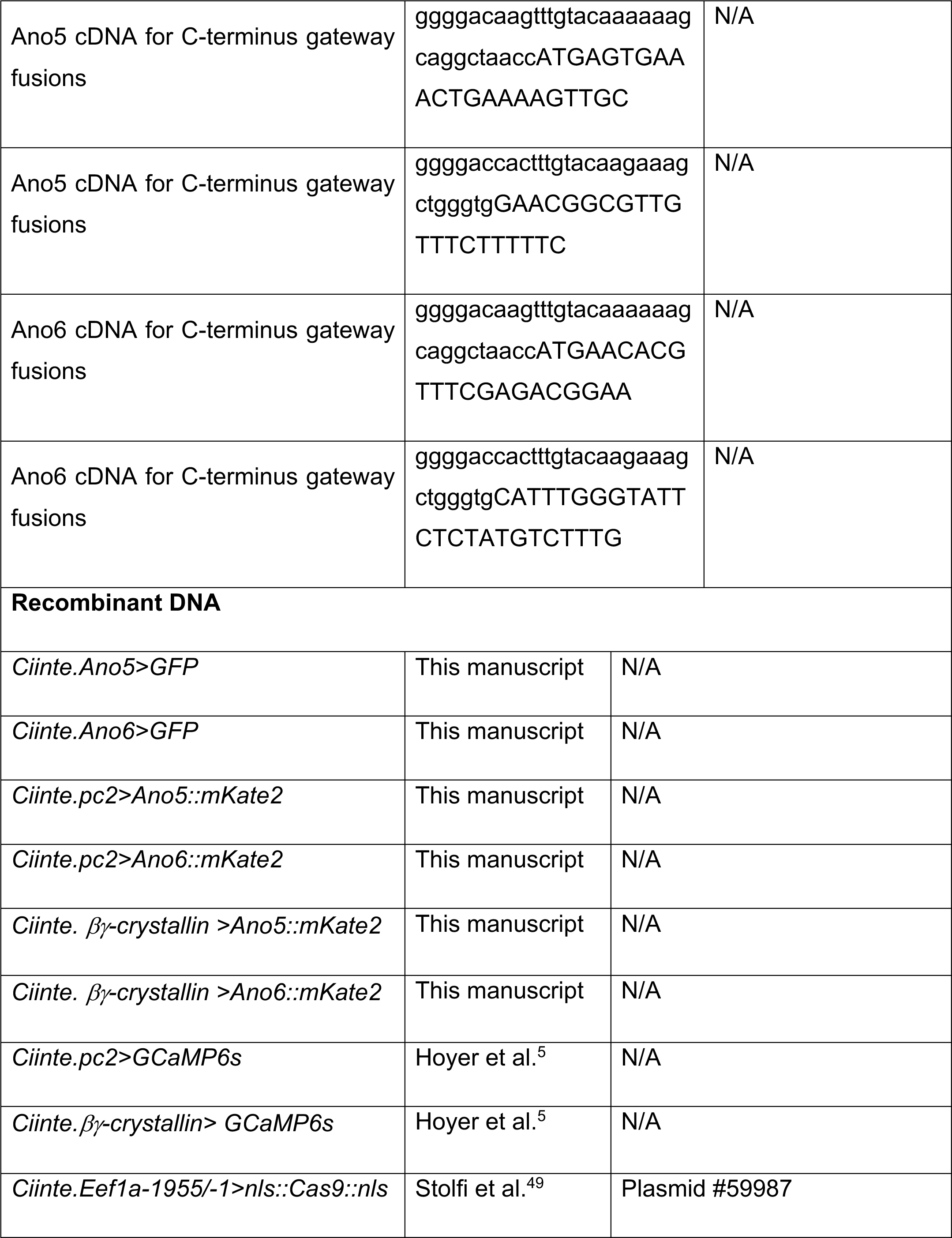
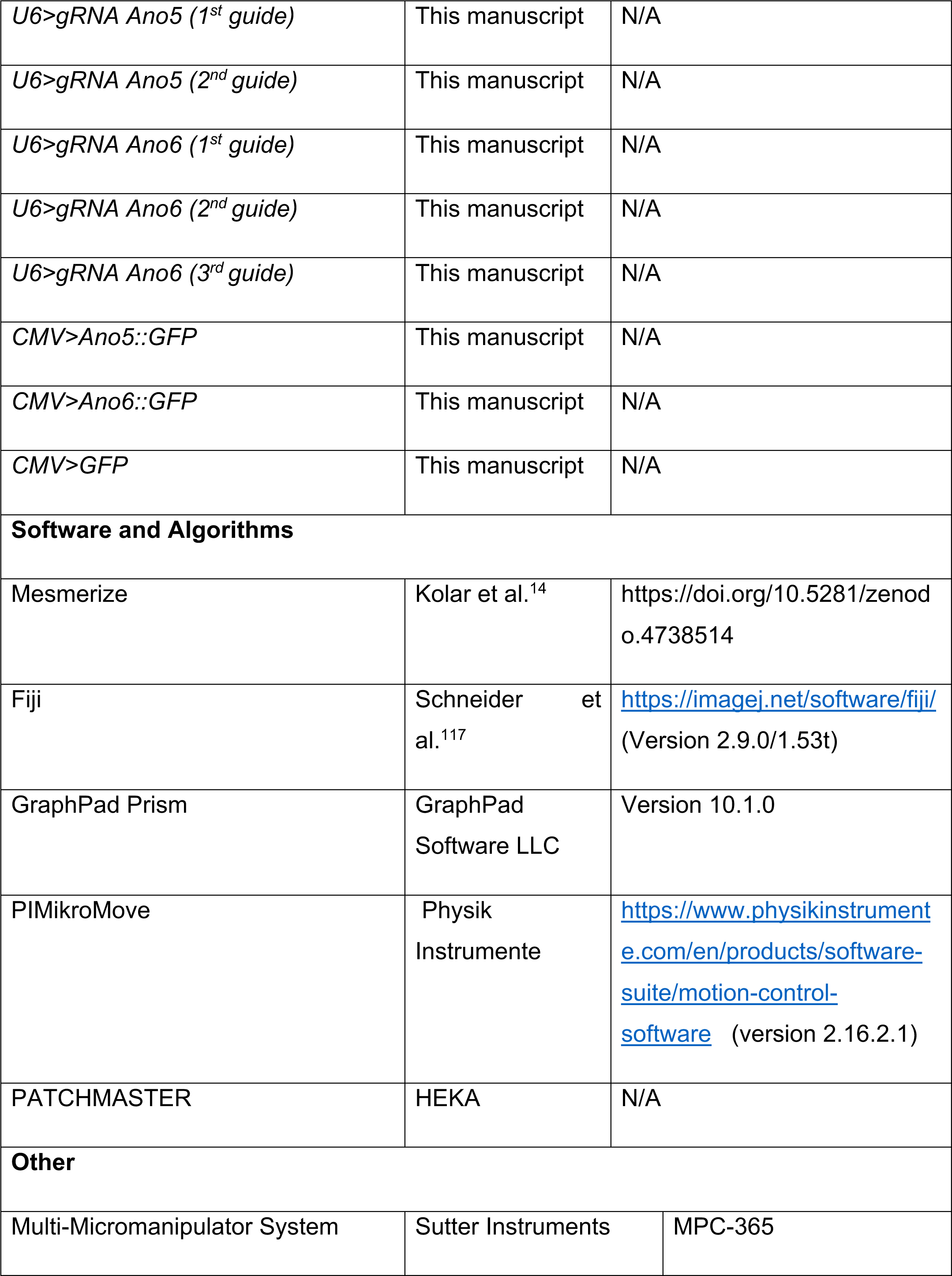
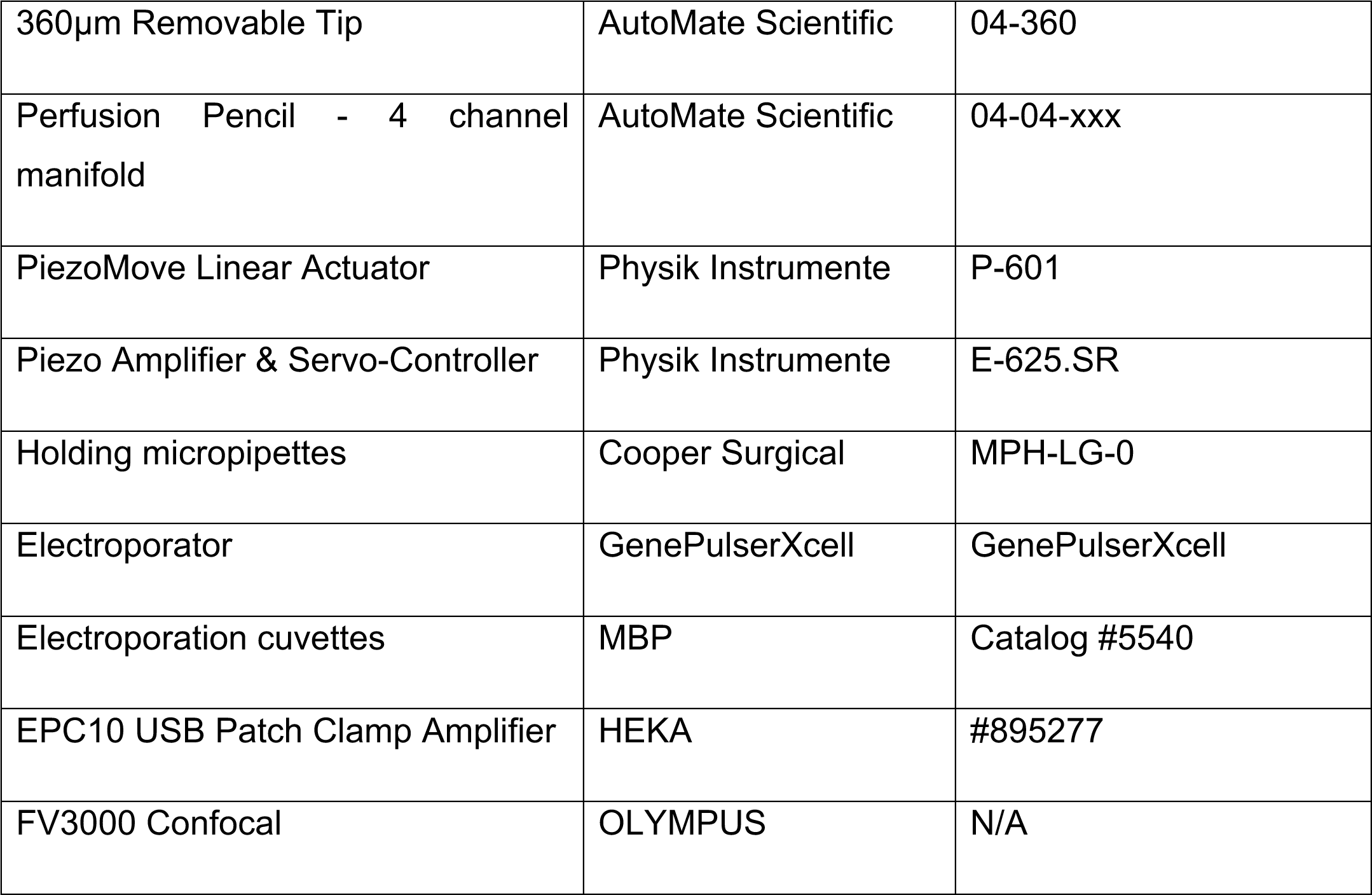

### Resource availability

#### Lead contact

Further information and requests for resources and reagents should be directed to and will be fulfilled by the lead contact, Marios Chatzigeorgiou

#### Materials availability

Plasmids generated in this study are available upon request from the lead contact.

### Data and code availability

#### Experimental model and study participant details

Adult *C. intestinalis* (previously known as Ciona intestinalis Type B) were sampled from: Døsjevika, Bildøy Marina AS, postcode 5353, in Bergen, Norway. The collection site can be located using the GPS coordinates: 60.344330, 5.110812. Adult *Cionas* were housed in our purpose-made facility at the Michael Sars Centre. <100 adults were kept in large 50L tanks with continuous sea water flow. We maintained a constant sea water temperature of 10°C with constant light and food supply (which incorporated a total of 4 species of diatoms and brown algae) to enhance egg production and reduce the propability of spontaneous spawning^50^.

### Method details

#### Electroporations of zygotes and staging of embryos

We dissected adult *C. intestinalis* to collect eggs and sperm for *in vitro* fertilization. We performed egg dechorionation using a 1% sodium thioglycolate and 0.1% pronase mix dissolved in filtered seawater. Eggs were mechanically agitated in dechorionation solution on a rocker for ca 10 min until we observed that they were dechorionated. We washed dechorionated eggs several times using ASW and then we fertilized with sperm for ∼10 min. After thoroughly washing the fertilized eggs, they were electroporated in a mix of mannitol solution (0.96M) with DNA ranging from 50 to 150μg. We electroporated using 4 mm gap electroporation cuvettes. Pulses were delivered using a BIORAD GenePulserXcell equipped with a CE-module. We used the subsequent parameters: Exponential Protocol: 50 V, Capacitance: 1000–1300μF and Resistance: ∞. We obtained electroporation time constants ranging from 10t to 20 milliseconds. We cultured embryos in ASW at 14°C. We adjusted the pH of the ASW to 8.4 at 14°C. The salinity of the ASW was set to 3.3–3.4%.

#### Molecular cloning

To amplify the promoters for Ano5 (KH.C8.274) and Ano6 (KH.C10.524) we used the *Ciona intestinalis* genome database Aniseed^51,52^ to obtain the cis-regulatory regions upstream of the two genes. Using Phusion-HF polymerase we obtained PCR products which were purified and inserted into the entry clone vector, pDONRP4-P1R using BP Clonase II (Invitrogen). The full length of the cDNA of Ano5, Ano6, mCh-Sec61 beta, and GPI-GFP were amplified by Phusion-HF polymerase based PCR and inserted into the entry clone vector, PDONR221 by BP Clonase II. The templates we used were local *Ciona* cDNA and mCh-Sec61 beta (mCh-Sec61 beta was a gift from Gia Voeltz (Addgene plasmid # 49155; http://n2t.net/addgene:49155; RRID:Addgene_49155) and pCAG:GPI-GFP (pCAG:GPI-GFP was a gift from Anna-Katerina Hadjantonakis (Addgene plasmid # 32601; http://n2t.net/addgene:32601; RRID:Addgene_32601). The expression plasmids were generated by LR reaction which recombined all the parts together into a pDESTII vector using the LR Clonase II enzyme (Invitrogen). The reporter proteins we used were UNC-76::GFP, GCaMP6s, mKate2, which were also cloned into the gateway system. For the mammalian cell expression plasmids, we amplified the cDNA of Ciona Ano5 and Ano6, with primers containing the ApaI and XhoI restriction sites which were used for subcloning of the cDNAs into mammalian cell line expression vectors.

#### Generation of mutant embryos by CRISPR/Cas9

The guide RNA(gRNA) sequences targeting *Ano5* and *Ano6* were designed through online tools E-CRISP http://www.e-crisp.org/E-CRISP/^53^ and CRISPOR (http://crispor.tefor.net/)^54^. Multiple pairs of gRNA were generated for each gene. The gRNA oligonucleotides were inserted into the U6 vector as described by Stolfi et al. ^49^. The control gRNA was designed to not target any *Ciona* genome by sequence blasting. 30μg EF1a>NLS::Cas9::NLS and 70μg of each U6>gRNA plasmid were electroporated together into the embryos. We used two gRNAs for *Ano5* and three gRNAs for *Ano6* to generate a deletion in genomic DNA. After reaching the larval stage, animals were transferred into a single tube of 8-Strip PCR Tubes. Single animal genomic DNA was extracted for genotyping and mutation detection. The primers used to amplify the regions harboring the target sites of the gRNAs for each gene were designed to cover all gRNA sites.

#### Metamorphosis assay

Around 8 hours after hatching, the larvae were transferred to a new 3cm petri dish without agarose coating. The petri dishes were filled with ASW with or without 10 mM NH_4_Cl solution. The animals were mounted on our Ciona Tracker 2.0 setups^16^, inside an incubator that kept the temperature constant at 14 °C. The larvae were illuminated using infrared LEDs (IR, peak emission 850 nm) and videos were recorded using the IR sensitive monochrome DMK 33UP1300 (Imaging Source) camera coupled to an MVL75M1 lens. We used custom made software to record 1 frame per second (60 frames per minute) for a maximum of 1740 minutes (29 hours). The software controlling the Ciona Tracker 2.0 and the video acquisition is available on GitHub: https://github.com/ChatzigeorgiouGroup/immobilize. Videos were analyzed using the markerless pose estimation software DeepLabCut^17^. Training of the model was done on 300 frames, from 5 videos, with 200 000 training iterations. We marked 6 points along the larvae (Point 1: Trunk tip/palps; Point 2: Trunk/Tail interface point; Points 3-5: equally spaced points along the tail Anterior to Posterior; Point 6: Tail tip). To quantify the rate of metamorphosis we measured the normalized length of the larvae over time (distance from point 1 trunk tip to point 6 tail tip).

#### Calcium imaging

A Zeiss Inverted microscope (ZEISS Axio Scope.A1) with a 40x water immersion lens was used for the poking and 10mM NH_4_Cl perfusion experiments. GCaMP6s expressing larvae were illuminated by a mercury lamp with a BP470/20, FT493, BP505-530 filter-set. We used a Hamamatsu Orca FlashV4 CMOS camera to acquire images at 10 Hz and exposure time 100ms. A custom made software was used for acquiring the images^55^. Stage 27 larvae electroporated with 80-100μg of either *Ciinte.pc2>GCaMP6s* or *Ciinte.βγ-crystallin>GCaMP6s* in combination with the plasmids for the genome editing (Cas9 and gRNAs) were screened for expression under a fluorescent stereomicroscope and transferred on an agarose-coated petri-dish with 0.03% MS-222. The target larva was sucked to a glass holding pipette at the neck-tail junction region by applying negative pressure provided by a 1ml syringe.

For the mechanical poke assay we recorded for a pre-stimulus period of 10 seconds. Upon the completion of this phase, a blunt glass needle attached to a micromanipulator (Sutter Instrument MPC-365 System) was moved automatically in the direction of the animal so that it would indent the pad of the papillae transiently before it was automatically drawn back to its starting position. The mean depression of the papillae from the poke stimulus delivery was 5,85 μm. The mean duration of the stimulus was 217ms. Following the end of the stimulus we continued recording calcium activity in the papillae for another 60 seconds, until the calcium signal had decreased to pre-stimulation levels.

For the chemical stimulation experiments the perfusion pencil which was used for ASW and 10mM NH_4_Cl perfusion was positioned at ca 250μm from the animal, pointing towards the papillae of the larvae. The first 10 seconds of the experiments the animals were perfused with a control solution consisting of ASW before switching to the stimulation solution for 10 seconds. We then switched back to perfusing with the control solution and kept recording the calcium activity in the papilla cells for a minimum post stimulation period of 60 seconds minute until the Ca^2+^ signal had returned to baseline levels. Data analysis was done in the Ca^2+^ imaging analysis platform Mesmerize^56^.

#### Cell culture and transfection

HEK293T cells were cultured in DMEM (Gibco^TM^), which was supplemented with 10% fetal bovine serum, 2 mM L-glutamine, 100 U/ml^-1^ penicillin, and 100 μg/ml^-1^ streptomycin. All transfections were carried out by Lipofectamine^TM^ Transfection Reagent (Invitrogen) following the manufacturer’s protocol. All experiments were performed between 36-48 h after the transfection.

#### Electrophysiology

Electrophysiological studies were carried out with the HEK293T cell after transfection from 36 to 48 hours. The cells were seeded on a round coverslip coated by poly l-lysine. The positive fluorescence cells were selected under microscopy (Zeiss). Recordings were performed using the whole-cell patch-clamp configuration. Patch pipettes had resistances ranged from 3–6 MΩ when filled with the pipette solution. Data were acquired by a HEKA EPC10 amplifier and Patchmaster software (HEKA). Current measurements were performed at room temperature. Capacitance and access resistance were monitored continuously. Currents were filtered at 2.9 kHz with a low-pass Bessel filter. The Pipette (intracellular) solution was composed of (mmol/L): 146 CsCl, 5 EGTA, 2 MgCl_2_, 10 sucrose, 10 HEPES, PH was adjusted to 7.3 with NMDG. The extracellular solution contained: 140mM NaCl, 5mM KCl, 2mM CaCl_2_, 1mM MgCl2, 10 mM HEPES, 15mM D-glucose, pH was adjusted to 7.4 with NaOH. Data were analyzed by python 3.8.

#### Scramblase assay

24 hours after transfection with Ano5::GFP, Ano6::GFP or control plasmids, the HEK293T cells were seeded in confocal imaging chamber. The cells were washed twice with PBS, then were incubated with 1:100 Alexa Fluor^TM^ 568 in binding buffer (Thermofisher) for 10 minutes, which also contained 1x RedDot™1 Far-Red Nuclear Stain (Biotium) for dead cell detection and with 10μM ionomycin or without for control. Then fixed the cells for 15 minutes with 4% Paraformaldehyde (PFA). The cells were washed three times then attached to a glass slide for imaging. The data was analyzed by Python 3.8 with segmentation to extract the green (GFP), red (Annexin v) and magenta (RedDot) signal information with manual correction by ImageJ. The cell numbers were counted by Scikit-image in python 3.8.

## Acknowledgments

We acknowledge funding from the Research Council of Norway: 339399 to M.C. and 234817 to the Michael Sars Centre.

## Author contributions

Z.L. and M.C. conceived the project. Z.L., J.H. and M.C. carried out the experiments. Z.L., J.H. analyzed the data. M.C. wrote the paper with input from all authors.

## References

1. Galindo, B.E., and Vacquier, V.D. (2005). Phylogeny of the TMEM16 protein family: Some members are overexpressed in cancer. International Journal of Molecular Medicine 16, 919–924.

2. Milenkovic, V.M., Brockmann, M., Stohr, H., Weber, B.H., and Strauss, O. (2010). Evolution and functional divergence of the anoctamin family of membrane proteins. BMC Evol Biol 10, 319. 10.1186/1471-2148-10-319.

3. Agostinelli, E., and Tammaro, P. (2022). Polymodal Control of TMEM16x Channels and Scramblases. Int J Mol Sci 23. 10.3390/ijms23031580.

4. Kalienkova, V., Clerico Mosina, V., and Paulino, C. (2021). The Groovy TMEM16 Family: Molecular Mechanisms of Lipid Scrambling and Ion Conduction. J Mol Biol 433, 166941. 10.1016/j.jmb.2021.166941.

5. Hoyer, J., Kolar, K., Athira, A., van den Burgh, M., Dondorp, D., Liang, Z., and Chatzigeorgiou, M. (2024). Polymodal sensory perception drives settlement and metamorphosis of Ciona larvae. Curr Biol 34, 1168–1182 e1167. 10.1016/j.cub.2024.01.041.

6. Johnson, C.J., Razy-Krajka, F., Zeng, F., Piekarz, K.M., Biliya, S., Rothbacher, U., and Stolfi, A. (2024). Specification of distinct cell types in a sensory-adhesive organ important for metamorphosis in tunicate larvae. PLoS Biol 22, e3002555. 10.1371/journal.pbio.3002555.

7. Dolcemascolo, G., Pennati, R., De Bernardi, F., Damiani, F., and Gianguzza, M. (2009). Ultrastructural comparative analysis on the adhesive papillae of the swimming larvae of three ascidian species. Isj-Invert Surviv J 6, S77–S86.

8. Pennati, R., Groppelli, S., De Bernardi, F., Mastrototaro, F., and Zega, G. (2009). Immunohistochemical analysis of adhesive papillae of Clavelina lepadiformis (Muller, 1776) and Clavelina phlegraea (Salfi, 1929) (Tunicata, Ascidiacea). Eur J Histochem 53, 25–33.

9. Zeng, F., Wunderer, J., Salvenmoser, W., Hess, M.W., Ladurner, P., and Rothbacher, U. (2018). Papillae revisited and the nature of the adhesive secreting collocytes. Dev Biol. 10.1016/j.ydbio.2018.11.012.

10. Liang, Z., Dondorp, D.C., and Chatzigeorgiou, M. (2023). Anoctamin 10/TMEM16K mediates convergent extension and tubulogenesis during notochord formation in the early chordate <em>Ciona intestinalis</em>. bioRxiv, 2023.2001.2020.524945. 10.1101/2023.01.20.524945.

11. Nakayama-Ishimura, A., Chambon, J.P., Horie, T., Satoh, N., and Sasakura, Y. (2009). Delineating metamorphic pathways in the ascidian Ciona intestinalis. Dev Biol 326, 357–367. 10.1016/j.ydbio.2008.11.026.

12. Torrence, S.A., and Cloney, R.A. (1983). Ascidian Larval Nervous-System - Primary Sensory Neurons in Adhesive Papillae. Zoomorphology 102, 111–123. Doi 10.1007/Bf00363804.

13. Cloney, R.A. (1979). Larval Adhesive Organs and Metamorphosis in Ascidians .2. Mechanism of Eversion of the Papillae of Distaplia-Occidentalis. Cell and Tissue Research 200, 453–473.

14. Cloney, R.A. (1977). Larval Adhesive Organs and Metamorphosis in Ascidians .1. Fine-Structure of Everting Papillae of Distaplia-Occidentalis. Cell and Tissue Research 183, 423–444.

15. Matsunobu, S., and Sasakura, Y. (2015). Time course for tail regression during metamorphosis of the ascidian Ciona intestinalis. Dev Biol 405, 71–81. 10.1016/j.ydbio.2015.06.016.

16. Athira, A., Dondorp, D., Rudolf, J., Peytral, O., and Chatzigeorgiou, M. (2022). Comprehensive analysis of locomotion dynamics in the protochordate Ciona intestinalis reveals how neuromodulators flexibly shape its behavioral repertoire. PLoS Biol 20, e3001744. 10.1371/journal.pbio.3001744.

17. Mathis, A., Mamidanna, P., Cury, K.M., Abe, T., Murthy, V.N., Mathis, M.W., and Bethge, M. (2018). DeepLabCut: markerless pose estimation of user-defined body parts with deep learning. Nat Neurosci 21, 1281–1289. 10.1038/s41593-018-0209-y.

18. Pedemonte, N., and Galietta, L.J. (2014). Structure and function of TMEM16 proteins (anoctamins). Physiol Rev 94, 419–459. 10.1152/physrev.00039.2011.

19. Meinertzhagen, I.A., Lemaire, P., and Okamura, Y. (2004). The neurobiology of the ascidian tadpole larva: recent developments in an ancient chordate. Annu Rev Neurosci 27, 453–485. 10.1146/annurev.neuro.27.070203.144255.

20. Horie, R., Hazbun, A., Chen, K., Cao, C., Levine, M., and Horie, T. (2018). Shared evolutionary origin of vertebrate neural crest and cranial placodes. Nature 560, 228–232. 10.1038/s41586-018-0385-7.

21. Patthey, C., Schlosser, G., and Shimeld, S.M. (2014). The evolutionary history of vertebrate cranial placodes--I: cell type evolution. Dev Biol 389, 82–97. 10.1016/j.ydbio.2014.01.017.

22. Cao, C., Lemaire, L.A., Wang, W., Yoon, P.H., Choi, Y.A., Parsons, L.R., Matese, J.C., Wang, W., Levine, M., and Chen, K. (2019). Comprehensive single-cell transcriptome lineages of a proto-vertebrate. Nature 571, 349–354. 10.1038/s41586-019-1385-y.

23. Mazet, F., Hutt, J.A., Milloz, J., Millard, J., Graham, A., and Shimeld, S.M. (2005). Molecular evidence from Ciona intestinalis for the evolutionary origin of vertebrate sensory placodes. Dev Biol 282, 494–508. 10.1016/j.ydbio.2005.02.021.

24. Liu, B., and Satou, Y. (2019). Foxg specifies sensory neurons in the anterior neural plate border of the ascidian embryo. Nat Commun 10, 4911. 10.1038/s41467-019-12839-6.

25. Wagner, E., Stolfi, A., Gi Choi, Y., and Levine, M. (2014). Islet is a key determinant of ascidian palp morphogenesis. Development 141, 3084–3092. 10.1242/dev.110684.

26. Poncelet, G., and Shimeld, S.M. (2020). The evolutionary origins of the vertebrate olfactory system. Open Biol 10, 200330. 10.1098/rsob.200330.

27. Kim, H., Kim, H., Nguyen, L.T., Ha, T., Lim, S., Kim, K., Kim, S.H., Han, K., Hyeon, S.J., Ryu, H., et al. (2022). Amplification of olfactory signals by Anoctamin 9 is important for mammalian olfaction. Prog Neurobiol 219, 102369. 10.1016/j.pneurobio.2022.102369.

28. Stephan, A.B., Shum, E.Y., Hirsh, S., Cygnar, K.D., Reisert, J., and Zhao, H.Q. (2009). ANO2 is the cilial calcium-activated chloride channel that may mediate olfactory amplification. P Natl Acad Sci USA 106, 11776–11781. 10.1073/pnas.0903304106.

29. Pifferi, S., Cenedese, V., and Menini, A. (2012). Anoctamin 2/TMEM16B: a calcium-activated chloride channel in olfactory transduction. Exp Physiol 97, 193–199. 10.1113/expphysiol.2011.058230.

30. Dibattista, M., Pifferi, S., Boccaccio, A., Menini, A., and Reisert, J. (2017). The long tale of the calcium activated Cl(-) channels in olfactory transduction. Channels (Austin) 11, 399–414. 10.1080/19336950.2017.1307489.

31. Pietra, G., Dibattista, M., Menini, A., Reisert, J., and Boccaccio, A. (2016). The Ca2+-activated Cl-channel TMEM16B regulates action potential firing and axonal targeting in olfactory sensory neurons. J Gen Physiol 148, 293–311. 10.1085/jgp.201611622.

32. Sakamoto, A., Hozumi, A., Shiraishi, A., Satake, H., Horie, T., and Sasakura, Y. (2022). The TRP channel PKD2 is involved in sensing the mechanical stimulus of adhesion for initiating metamorphosis in the chordate Ciona. Dev Growth Differ 64, 395–408. 10.1111/dgd.12801.

33. Guarneri, G., Pifferi, S., Dibattista, M., Reisert, J., and Menini, A. (2023). Paradoxical electro-olfactogram responses in TMEM16B knock-out mice. Chem Senses 48. 10.1093/chemse/bjad003.

34. Huang, W.C., Xiao, S., Huang, F., Harfe, B.D., Jan, Y.N., and Jan, L.Y. (2012). Calcium-activated chloride channels (CaCCs) regulate action potential and synaptic response in hippocampal neurons. Neuron 74, 179–192. 10.1016/j.neuron.2012.01.033.

35. Ha, G.E., Lee, J., Kwak, H., Song, K., Kwon, J., Jung, S.Y., Hong, J., Chang, G.E., Hwang, E.M., Shin, H.S., et al. (2016). The Ca(2+)-activated chloride channel anoctamin-2 mediates spike-frequency adaptation and regulates sensory transmission in thalamocortical neurons. Nat Commun 7, 13791. 10.1038/ncomms13791.

36. Wang, L., Simms, J., Peters, C.J., Tynan-La Fontaine, M., Li, K., Gill, T.M., Jan, Y.N., and Jan, L.Y. (2019). TMEM16B Calcium-Activated Chloride Channels Regulate Action Potential Firing in Lateral Septum and Aggression in Male Mice. J Neurosci 39, 7102–7117. 10.1523/JNEUROSCI.3137-18.2019.

37. Li, K.X., He, M., Ye, W., Simms, J., Gill, M., Xiang, X., Jan, Y.N., and Jan, L.Y. (2019). TMEM16B regulates anxiety-related behavior and GABAergic neuronal signaling in the central lateral amygdala. Elife 8. 10.7554/eLife.47106.

38. Zak, J.D., Grimaud, J., Li, R.C., Lin, C.C., and Murthy, V.N. (2018). Calcium-activated chloride channels clamp odor-evoked spike activity in olfactory receptor neurons. Sci Rep 8, 10600. 10.1038/s41598-018-28855-3.

39. Henkel, B., Drose, D.R., Ackels, T., Oberland, S., Spehr, M., and Neuhaus, E.M. (2015). Co-expression of anoctamins in cilia of olfactory sensory neurons. Chem Senses 40, 73–87. 10.1093/chemse/bju061.

40. Yang, H., Kim, A., David, T., Palmer, D., Jin, T., Tien, J., Huang, F., Cheng, T., Coughlin, S.R., Jan, Y.N., and Jan, L.Y. (2012). TMEM16F forms a Ca2+-activated cation channel required for lipid scrambling in platelets during blood coagulation. Cell 151, 111–122. 10.1016/j.cell.2012.07.036.

41. Grubb, S., Poulsen, K.A., Juul, C.A., Kyed, T., Klausen, T.K., Larsen, E.H., and Hoffmann, E.K. (2013). TMEM16F (Anoctamin 6), an anion channel of delayed Ca(2+) activation. J Gen Physiol 141, 585–600. 10.1085/jgp.201210861.

42. Shimizu, T., Iehara, T., Sato, K., Fujii, T., Sakai, H., and Okada, Y. (2013). TMEM16F is a component of a Ca2+-activated Cl-channel but not a volume-sensitive outwardly rectifying Cl-channel. Am J Physiol Cell Physiol 304, C748–759. 10.1152/ajpcell.00228.2012.

43. Scudieri, P., Caci, E., Venturini, A., Sondo, E., Pianigiani, G., Marchetti, C., Ravazzolo, R., Pagani, F., and Galietta, L.J.V. (2015). Ion channel and lipid scramblase activity associated with expression of TMEM16F/ANO6 isoforms. The Journal of Physiology 593, 3829–3848. 10.1113/JP270691.

44. Tian, Y., Schreiber, R., and Kunzelmann, K. (2012). Anoctamins are a family of Ca2+-activated Cl-channels. J Cell Sci 125, 4991–4998. 10.1242/jcs.109553.

45. Bushell, S.R., Pike, A.C.W., Falzone, M.E., Rorsman, N.J.G., Ta, C.M., Corey, R.A., Newport, T.D., Christianson, J.C., Scofano, L.F., Shintre, C.A., et al. (2019). The structural basis of lipid scrambling and inactivation in the endoplasmic reticulum scramblase TMEM16K. Nat Commun 10, 3956. 10.1038/s41467-019-11753-1.

46. Malvezzi, M., Chalat, M., Janjusevic, R., Picollo, A., Terashima, H., Menon, A.K., and Accardi, A. (2013). Ca2+-dependent phospholipid scrambling by a reconstituted TMEM16 ion channel. Nat Commun 4, 2367. 10.1038/ncomms3367.

47. Lee, B.C., Menon, A.K., and Accardi, A. (2016). The nhTMEM16 Scramblase Is Also a Nonselective Ion Channel. Biophys J 111, 1919–1924. 10.1016/j.bpj.2016.09.032.

48. Jiang, T., Yu, K., Hartzell, H.C., and Tajkhorshid, E. (2017). Lipids and ions traverse the membrane by the same physical pathway in the nhTMEM16 scramblase. Elife 6. 10.7554/eLife.28671.

49. Stolfi, A., Gandhi, S., Salek, F., and Christiaen, L. (2014). Tissue-specific genome editing in Ciona embryos by CRISPR/Cas9. Development 141, 4115–4120. 10.1242/dev.114488.

50. Rudolf, J., Dondorp, D., Canon, L., Tieo, S., and Chatzigeorgiou, M. (2019). Automated behavioural analysis reveals the basic behavioural repertoire of the urochordate Ciona intestinalis. Sci Rep 9, 2416. 10.1038/s41598-019-38791-5.

51. Dardaillon, J., Dauga, D., Simion, P., Faure, E., Onuma, T.A., DeBiasse, M.B., Louis, A., Nitta, K.R., Naville, M., Besnardeau, L., et al. (2019). ANISEED 2019: 4D exploration of genetic data for an extended range of tunicates. Nucleic Acids Res. 10.1093/nar/gkz955.

52. Brozovic, M., Dantec, C., Dardaillon, J., Dauga, D., Faure, E., Gineste, M., Louis, A., Naville, M., Nitta, K.R., Piette, J., et al. (2018). ANISEED 2017: extending the integrated ascidian database to the exploration and evolutionary comparison of genome-scale datasets. Nucleic Acids Res 46, D718–D725. 10.1093/nar/gkx1108.

53. Heigwer, F., Kerr, G., and Boutros, M. (2014). E-CRISP: fast CRISPR target site identification. Nature Methods 11, 122–123. 10.1038/nmeth.2812.

54. Concordet, J.P., and Haeussler, M. (2018). CRISPOR: intuitive guide selection for CRISPR/Cas9 genome editing experiments and screens. Nucleic Acids Res 46, W242–W245. 10.1093/nar/gky354.

55. Kolar, K., and Chatzigeorgiou, M. (2019). Simple GUI for acquiring images from a Hamamatsu Orca Flash 4.0 CMOS camera. Zenodo. 10.5281/zenodo.3370464.

56. Kolar, K., Dondorp, D., Zwiggelaar, J.C., Høyer, J., and Chatzigeorgiou, M. (2021). Mesmerize is a dynamically adaptable user-friendly analysis platform for 2D and 3D calcium imaging data. Nature Communications 12. 10.1038/s41467-021-26550-y.

