## Supplementary material for "Anoctamins mediate polymodal sensory perception and larval metamorphosis in a non-vertebrate chordate": Data S1 Table Legends

### SUPPLEMENTAL TABLE LEGENDS

**Data S1 Statistical analyses and additional quantitative analyses. Related to STAR Methods, Figures 1-3, and Figures S1 and S2.**

A) Quantification of the frequency with which *Ciinte.Ano5>GFP* and *Ciinte.Ano6>GFP*, are expressed in different cell types of the larval nervous system. Related to Figure 1A, 1B and Figure S1A-S1G.

B) Kruskal-Wallis test followed by Dunn's multiple comparisons test of larval normalized body length between negative control, $Ano5^{CRISPR}$, $Ano6^{CRISPR}$ and $Ano5^{CRISPR}$; $Ano6^{CRISPR}$ larvae at 10 hours in ASW, related to Figure 1F, Figure S2A.

C) Kruskal-Wallis test followed by Dunn's multiple comparisons test of larval normalized body length between negative control, $Ano5^{CRISPR}$, $Ano6^{CRISPR}$ and $Ano5^{CRISPR}$; $Ano6^{CRISPR}$ larvae at 20 hours in ASW, related to Figure 1F, Figure S2B.

D) Kruskal-Wallis test followed by Dunn's multiple comparisons test of larval normalized body length between negative control, $Ano5^{CRISPR}$, $Ano6^{CRISPR}$ and $Ano5^{CRISPR}$; $Ano6^{CRISPR}$ larvae at 2 hours in 10mM $NH_4Cl$, related to Figure 1G, Figure S2C.

E) Kruskal-Wallis test followed by Dunn's multiple comparisons test of larval normalized body length between negative control, $Ano5^{CRISPR}$, $Ano6^{CRISPR}$ and $Ano5^{CRISPR}$; $Ano6^{CRISPR}$ larvae at 8 hours in 10mM $NH_4Cl$, related to Figure 1G, Figure S2D.

F) Kruskal-Wallis test followed by Dunn's multiple comparisons test of PSN $Ca2^+$ peak amplitude in response to mechanical poke between negative control, $Ano5^{CRISPR}$, $Ano6^{CRISPR}$ and $Ano5^{CRISPR}$; $Ano6^{CRISPR}$ larvae, related to Figure 2D, Figure S2E.

G) Kruskal-Wallis test followed by Dunn's multiple comparisons test of PSN $Ca2^+$ peak area in response to mechanical poke between negative control, $Ano5^{CRISPR}$, $Ano6^{CRISPR}$ and $Ano5^{CRISPR}$; $Ano6^{CRISPR}$ larvae, related to Figure 2D, Figure S2F.

H) Kruskal-Wallis test followed by Dunn's multiple comparisons test of PSN $Ca2^+$ peak rise time in response to mechanical poke between negative control, $Ano5^{CRISPR}$, $Ano6^{CRISPR}$ and $Ano5^{CRISPR}$; $Ano6^{CRISPR}$ larvae, related to Figure 2D, Figure S2G.

I) Kruskal-Wallis test followed by Dunn's multiple comparisons test of PSN $Ca2^+$ peak fall time in response to mechanical poke between negative control, $Ano5^{CRISPR}$, $Ano6^{CRISPR}$ and $Ano5^{CRISPR}$; $Ano6^{CRISPR}$ larvae, related to Figure 2D, Figure S2H.

J) Kruskal-Wallis test followed by Dunn's multiple comparisons test of PSN $Ca2^+$ peak duration in response to mechanical poke between negative control, $Ano5^{CRISPR}$, $Ano6^{CRISPR}$ and $Ano5^{CRISPR}$; $Ano6^{CRISPR}$ larvae, related to Figure 2D, Figure S2I.

K) Kruskal-Wallis test followed by Dunn's multiple comparisons test of PSN $Ca2^+$ peak amplitude in response to 10mM $NH_4Cl$ between negative control, $Ano5^{CRISPR}$, $Ano6^{CRISPR}$ and $Ano5^{CRISPR}$; $Ano6^{CRISPR}$ larvae, related to Figure 2F, Figure S2J.

L) Kruskal-Wallis test followed by Dunn's multiple comparisons test of PSN $Ca2^+$ peak area in response to 10mM $NH_4Cl$ between negative control, $Ano5^{CRISPR}$, $Ano6^{CRISPR}$ and $Ano5^{CRISPR}$; $Ano6^{CRISPR}$ larvae, related to Figure 2F, Figure S2K.

M) Kruskal-Wallis test followed by Dunn's multiple comparisons test of PSN $Ca2^+$ peak rise time in response to 10mM $NH_4Cl$ between negative control, $Ano5^{CRISPR}$, $Ano6^{CRISPR}$ and $Ano5^{CRISPR}$; $Ano6^{CRISPR}$ larvae, related to Figure 2F, Figure S2L.

N) Kruskal-Wallis test followed by Dunn's multiple comparisons test of PSN $Ca2^+$ peak fall time in response to 10mM $NH_4Cl$ between negative control, $Ano5^{CRISPR}$, $Ano6^{CRISPR}$ and $Ano5^{CRISPR}$; $Ano6^{CRISPR}$ larvae, related to Figure 2F, Figure S2M.

O) Kruskal-Wallis test followed by Dunn's multiple comparisons test of PSN $Ca2^+$ peak duration in response to 10mM $NH_4Cl$ between negative control, $Ano5^{CRISPR}$, $Ano6^{CRISPR}$ and $Ano5^{CRISPR}$; $Ano6^{CRISPR}$ larvae, related to Figure 2F, Figure S2N.

P) Kruskal-Wallis test followed by Dunn's multiple comparisons test of ACC $Ca2^+$ peak amplitude in response to mechanical poke between negative control, $Ano5^{CRISPR}$, $Ano6^{CRISPR}$ and $Ano5^{CRISPR}$; $Ano6^{CRISPR}$ larvae, related to Figure 2J, Figure S2O.

Q) Kruskal-Wallis test followed by Dunn's multiple comparisons test of ACC $Ca2^+$ peak area in response to mechanical poke between negative control, $Ano5^{CRISPR}$, $Ano6^{CRISPR}$ and $Ano5^{CRISPR}$; $Ano6^{CRISPR}$ larvae, related to Figure 2J, Figure S2P.

R) Kruskal-Wallis test followed by Dunn's multiple comparisons test of ACC $Ca2^+$ peak rise time in response to mechanical poke between negative control, $Ano5^{CRISPR}$, $Ano6^{CRISPR}$ and $Ano5^{CRISPR}$; $Ano6^{CRISPR}$ larvae, related to Figure 2J, Figure S2Q.

S) Kruskal-Wallis test followed by Dunn's multiple comparisons test of ACC $Ca2^+$ peak fall time in response to mechanical poke between negative control, $Ano5^{CRISPR}$, $Ano6^{CRISPR}$ and $Ano5^{CRISPR}$; $Ano6^{CRISPR}$ larvae, related to Figure 2J, Figure S2R.

T) Kruskal-Wallis test followed by Dunn's multiple comparisons test of ACC $Ca2^+$ peak duration in response to mechanical poke between negative control, $Ano5^{CRISPR}$, $Ano6^{CRISPR}$ and $Ano5^{CRISPR}$; $Ano6^{CRISPR}$ larvae, related to Figure 2J, Figure S2S.

U) Kruskal-Wallis test followed by Dunn's multiple comparisons test of ACC $Ca2^+$ peak amplitude in response to 10mM $NH_4Cl$ between negative control, $Ano5^{CRISPR}$, $Ano6^{CRISPR}$ and $Ano5^{CRISPR}$; $Ano6^{CRISPR}$ larvae, related to Figure 2L, Figure S2T.

V) Kruskal-Wallis test followed by Dunn's multiple comparisons test of ACC $Ca2^+$ peak area in response to 10mM $NH_4Cl$ between negative control, $Ano5^{CRISPR}$, $Ano6^{CRISPR}$ and $Ano5^{CRISPR}$; $Ano6^{CRISPR}$ larvae, related to Figure 2L, Figure S2U.

W) Kruskal-Wallis test followed by Dunn's multiple comparisons test of ACC $Ca2^+$ peak rise time in response to 10mM $NH_4Cl$ between negative control, $Ano5^{CRISPR}$, $Ano6^{CRISPR}$ and $Ano5^{CRISPR}$; $Ano6^{CRISPR}$ larvae, related to Figure 2L, Figure S2V.

X) Kruskal-Wallis test followed by Dunn's multiple comparisons test of ACC $Ca2^+$ peak fall time in response to 10mM $NH_4Cl$ between negative control, $Ano5^{CRISPR}$, $Ano6^{CRISPR}$ and $Ano5^{CRISPR}$; $Ano6^{CRISPR}$ larvae, related to Figure 2L, Figure S2W.

Y) Kruskal-Wallis test followed by Dunn's multiple comparisons test of ACC $Ca2^+$ peak duration in response to 10mM $NH_4Cl$ between negative control, $Ano5^{CRISPR}$, $Ano6^{CRISPR}$ and $Ano5^{CRISPR}$; $Ano6^{CRISPR}$ larvae, related to Figure 2L, Figure S2X.

Z) Quantification of the heterelogous experiments to assay the scramblase activity of *Ciona intestinalis* Ano5 and Ano6, related to Figure 3A.
